# Expression levels and dimer abundance of lamin A/C direct nuclear shape integrity in malignant cancer cells

**DOI:** 10.64898/2026.05.11.724268

**Authors:** Myron Nicolaas Franciscus Hensgens, Aditya Mhaskar, Hylkje Geertsema

## Abstract

Abnormalities in nuclear morphology are an important diagnostic tool to determine malignancy in cancer cells and are characterised by nuclear blebbing and deformations. Nuclear shape is mostly maintained by a dense protein meshwork of lamins, consisting of 4 lamin subtypes, of which the individual contribution to nuclear shape maintenance remains elusive. In this study, we decouple the roles of lamin A, C, and B1 across cancer cell lines with varying malignant potential (HeLa, HT1080, and MDA-MB-231). Using single-cell correlation analysis, we directly link reduced lamin A/C, and not lamin B1, expression levels to nuclear deformability. We found that the nuclear shape of the more malignant MDA-MB-231 cells is approximately 4-fold more sensitive to lamin A/C than HeLa and HT1080 cells. Biochemical analyses reveal cell-type-specific variation in lamin A/C interactions and homodimer formation that correlates with nuclear shape deformations. In contrast to healthy mouse embryonic fibroblast cells, malignant cells exhibit reduced dimerisation, which correlates with nuclear deformability. As such, our study links, for the first time, the lamin A/C dimerisation state to nuclear abnormalities, thereby providing new avenues for investigating cancer progression.

## Introduction

Nuclear morphology has been widely employed as a diagnostic tool for cancer and metastatic prognosis.^1^ Abnormalities of the natural spherical and ellipsoidal nuclear shape have been correlated to a broad spectrum of diseases in metazoan cells, including cancer.^1,2^ Importantly, the more malignant cancer types have been related to severe nuclear deformations. Consequently, nuclear abnormalities supposedly relate to a mechanical shift towards a softer nucleus, which enables migration through confined spaces and thus enhances metastatic potential.^3^ In addition, nuclear deformations could facilitate biochemical signalling pathways that promote metastatic progression.^4^ As a consequence, maintaining a round nucleus is important for cellular health and is supported by a highly conserved protein meshwork that underlies the nuclear membrane, the nuclear lamina.

The nuclear lamina consists of type V intermediate filaments that are subcategorised into two classes, A-type lamins and B-type lamins. The A-type lamins are comprised of lamin A and its splice variant, lamin C, and have distinct hydrophilic and viscoelastic properties.^5^ The B-type lamins are encoded by separate genes, lamin B1 by LMNB1 and lamin B2 by LMNB2, and are hydrophobic and elastic.^6–9^ Both lamin types assemble into polymers and filaments, which subsequently form distinct, dense meshworks that range from 10 to 100 nm in thickness.^10–12^ The proportional combination of B-type (i.e. elastic) and A-type (i.e. stiff) lamins enables the nucleus to regulate its shape.^13–15^

Lower expression of A-type lamins has been associated with greater nuclear deformability and more invasive and aggressive cancer variants, whereas higher expression of A-type lamins has been associated with increased cancer survival and nuclear rigidity.^13,14,16–18^ Reduced lamin A expression is therefore believed to be the primary indicator of metastatic prognosis and cancer recurrence.^16–18^ Further, lamin B1 downregulation has been shown to alter nuclear morphology and result in a more malignant cancer phenotype in breast cancer cells.^4^ Nevertheless, the relationship between lamin expression and metastasis prognoses appears to be complex and tumour-type specific. Here, we aim to provide further insight into the contributions of individual lamin types to nuclear shape regulation across different cancer cell types.

In this study, we utilise single-cell correlation analysis across three wild-type human cancer cell lines, the cervical cancer cell line HeLa, the fibrosarcoma cell line HT1080 and the breast cancer cell line MDA-MB-231. By using fluorescence confocal microscopy on cells immunostained for lamin A/C and B1, we decouple the roles of endogenous lamin A/C and lamin B1 expression and their impact on nuclear shape, enabling a quantitative comparison of lamin expression levels and nuclear abnormalities. Our results demonstrate a clear dependence of lamin A/C, but not lamin B1, expression on nuclear deformability. At the same time, we also show cell-to-cell variability in the response to lamin A/C on the nuclear shape, reflecting the underlying complexity of nuclear envelope deformations and maintenance. Surprisingly, we have uncovered a correlation between lamin A/C dimerisation and cell-type specific nuclear deformability, suggesting that lamin A/C interaction strength, presumably facilitated by post-translational modifications, fulfils a novel and critical role on nuclear shape regulation.

## Results

### Population-averaged intensities of lamin A/C and B1 expression levels do not explain nuclear shape diversity between cell lines

We have employed 3D confocal fluorescence microscopy on HeLa, HT1080, MDA-MB-231, and NIH3T3 cells that were immunostained for lamin A/C and lamin B1. HeLa cells were chosen as a less invasive, well-studied cancer cell line, HT1080 and MDA-MB-231 cells were selected for their metastatic potential and are known to be highly aggressive and invasive.^19–23^ NIH3T3 cells were selected as a bona fide lamin B1-deficient cell line, although they are mouse cells compared to human cells.^24^

Initial qualitative observations revealed distinct differences in nuclear morphology across the cell lines (Figure 1A). We quantified these nuclear shape deformations by determining the nuclear solidity of individual cells, defined as the area of the fluorescent signal divided by the calculated area of the convex hull (Figure 1B). Exact overlap of the lamin signal with the convex hull, as in a perfect smooth elliptic nucleus, resulted in a solidity value of 1.0. On the contrary, folded or invaginated nuclei poorly align with the convex hull, leading to a solidity value below 1.0. Our results show that the aggressive human metastatic cancer cell lines had a significant reduction in their solidity, whereas the mouse embryonic fibroblast cell line showed very rounded nuclei, followed by the less invasive HeLa cells (Figure 1C).

**Figure 1.**
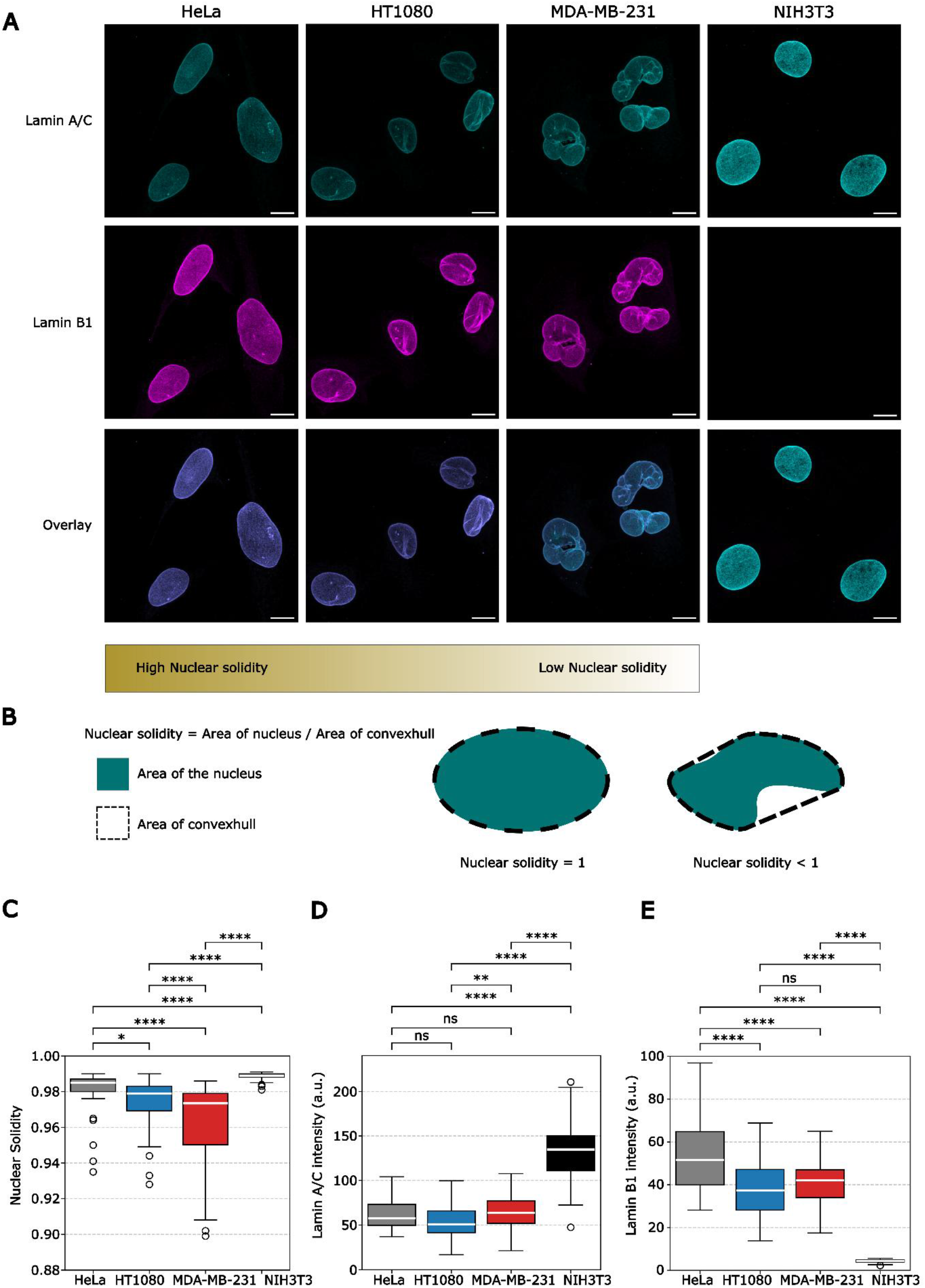
Comparison of lamin A/C and B1 expression and nuclear solidity across cell lines. **A**) Maximum-intensity projections of HeLa, HT1080, MDA-MB-231 and NIH3T3 cell lines immunostained for lamin A/C and B1 indicated NIH3T3 to comprise the most round nuclei, followed by HeLa, HT1080, and, lastly, MDA-MB-231, which showed the most disturbed nuclear shapes. NIH3T3 cells did not express lamin B1. **B**) Quantification of nuclear geometry was done by determining the nuclear solidity. Nuclear solidity is calculated by the detected area, isolated from the maximum-intensity projections, divided by the area of the convex hull. **C**) Boxplots of the obtained nuclear solidity for HeLa, HT1080, MDA-MB-231 and NIH3T3. **D**) Boxplots of the fluorescent intensity levels of lamin A/C for HeLa, HT1080, MDA-MB-231 and NIH3T3. **E**) Boxplots of the fluorescent intensity levels of lamin B1 for HeLa, HT1080, MDA-MB-231 and NIH3T3. Scale bars in **A**) represent 10 μm. P-values for the boxplots were determined by using a two-sided unpaired Student’s t-test. Data was assumed to be normally distributed with comparable variance. * are p-values<0.05, ** are p-values < 1*10^−2^, *** are p-values < 1*10^−3^, **** are p-values < 1*10^−4^ and not significant p-values are shown by ‘ns’. The number of nuclei imaged is n = 70 for HeLa cells, n = 82 for HT1080 cells, n = 64 for MDA-MB-231 cells, and n = 54 for NIH3T3 cells. Imaged over three consecutive experiments.

We next questioned how these different nuclear morphologies are supported by the expression levels of the different lamin types. To this end, we evaluated fluorescent intensities of lamin A/C and B1, measured with the same settings across all measurements, as a proxy for expression patterns across all cell lines. This assay showed that HeLa cells contain the highest amount of lamin B1 and have similar levels of lamin A/C as HT1080 and MDA-MB-231 (Figure 1D, E). Comparison of HT1080 and MDA-MB-231 reveal small, but significant, differences in lamin A/C expression but no difference in lamin B1 expression (Figure 1D, E). In contrast, NIH3T3 cells express almost no lamin B1, in agreement with previous findings, but show the highest level of lamin A/C compared with other cells (Figure 1D, E).^24^

Combining our results on the expression of lamin A/C and B1 and the cell’s nuclear solidity, however, did not show a clear correlation. NIH3T3 showed the most regularly shaped nuclei, but did not exhibit any lamin B1 signal, suggesting a key role of lamin A/C in maintaining a smooth, rounded nucleus. Though, HeLa cells exhibit the highest levels of lamin B1 and contain the second most regularly shaped nuclei, with occasional folds. In contrast, HeLa’s lamin A/C expression levels were similar to HT1080 and MDA-MB-231, although the HT1080 and MDA-MB-231 had more irregularly shaped nuclei, with the MDA-MB-231 containing the most abnormalities (Figure 1C, 1D). We hypothesised that the population-averaged nature of the pooled fluorescence data might mask the underlying relationship between lamin expression and nuclear deformability. Consequently, we performed single-cell correlation analysis of lamin expression levels and nuclear shape.

### Single-cell correlation analysis shows co-expression of lamin A/C and lamin B1

We first set out to examine whether nucleus deformations might be facilitated by either lower lamin A/C or lamin B1 expression levels, or both. To this end, we plotted the lamin A/C against lamin B1 fluorescence intensities for single cells, and included the nuclear solidity values in this plot (Figure 2). Interestingly, we found that all cell lines exhibited a consistent stoichiometric ratio of lamin A/C and B1, as lamin A/C and lamin B1 expression showed a significant linear correlation, indicating correlated expression of both lamin types. HeLa cells revealed the strongest co-expression as the fitted slope was 0.732, whereas the fitted slope was 0.248 for HT1080 cells, and 0.167 for MDA-MB-231 cells (Table 1). In comparison, HT1080 cells exhibited more nuclei with lower lamin B1 expression upon increased lamin A/C levels (Figure 2B). MDA-MB-231 cells showed the same trend towards lower lamin B1 expression but differed from the HT1080 cell line by lacking the subpopulation that expressed low lamin A/C and lamin B1 levels (Figure 2C). Consequently, MDA-MB-231 showed the highest variability in lamin A/C and lamin B1 stoichiometry and exposed the most lamin A/C-dominated phenotypes. Together, these observations reveal a higher degree of decoupling of lamin A/C and B1 expression for the more metastatic cancer cell lines. Representative images across all three human cancer cell lines (Figure 2A-C) further illustrate the co-expression of lamin subtypes. NIH3T3 was also investigated, but the very low lamin B1 expression levels opposed the observation of lamin B1 and lamin A/C co-expression (Sup. Fig. 1). The species difference and significantly altered lamin A/C and B1 expression patterns led us to focus more on the three human cancer cell lines.

**Table 1:** Statistical parameters of lamin A/C and B1 co-expression across cell lines.

|  |  |  |  |  |
| --- | --- | --- | --- | --- |
| Cell types: | Slope (B1:A/C) | Pearson coefficient r | Co-expression strength of B1:A/C | p-value |
| HeLa | 0.732 | 0.692 | High | p < 0.001 |
| HT1080 | 0.248 | 0.474 | Moderate | P < 0.001 |
| MDA-MB-231 | 0.167 | 0.336 | Low | P < 0.01 |

**Figure 2.**
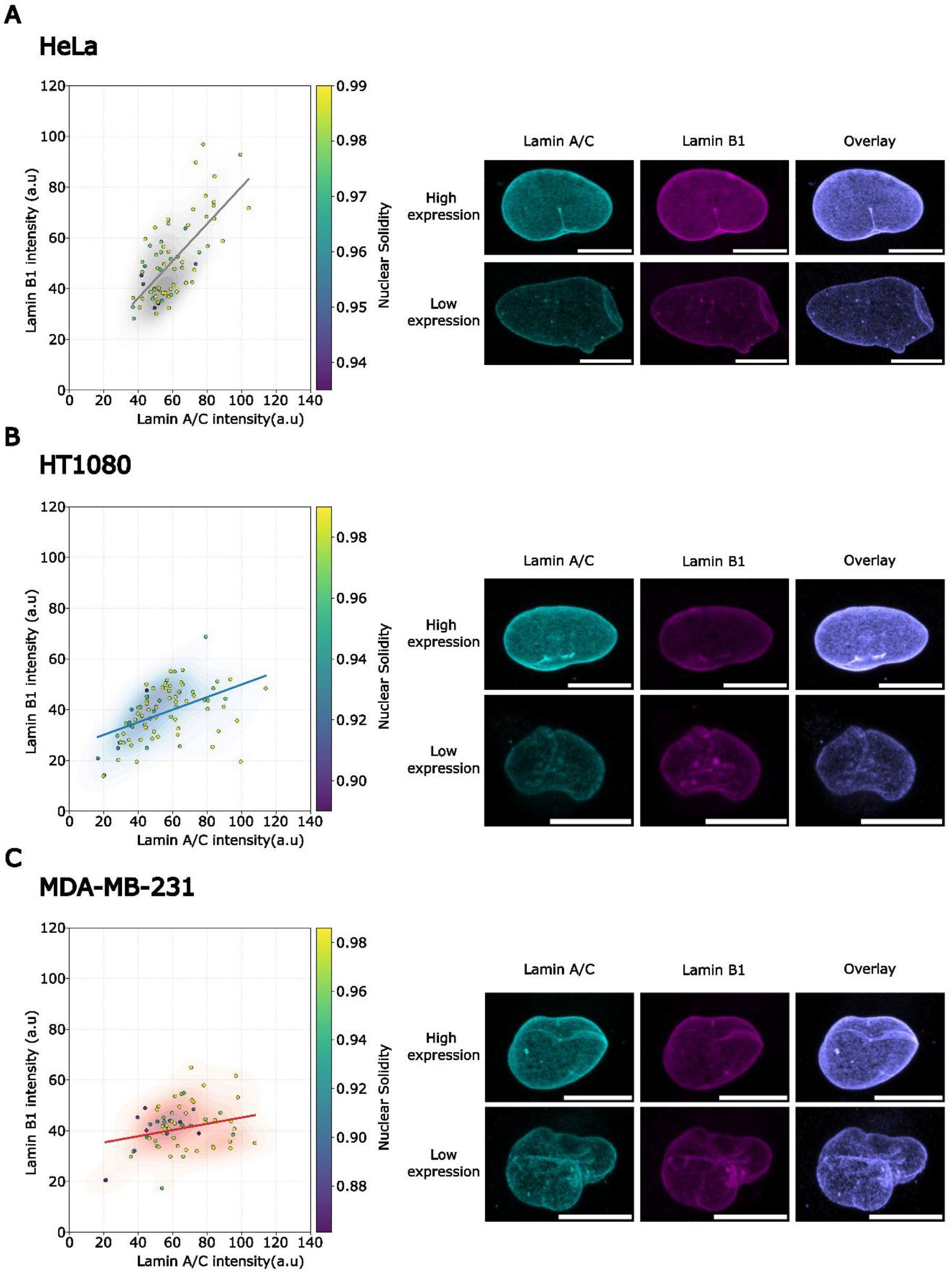
Single-cell analysis of lamin A/C and B1 co-expression reveals stoichiometric decoupling in metastatic cells. **A**) For HeLa cells, lamin A/C fluorescent intensities plotted against lamin B1 were fitted with a linear trendline. The trendline has a Pearson coefficient of 0.692 and a slope of 0.732. The data points are additionally colour-coded for the corresponding nuclear solidity. Representative images of HeLa nuclei expressing high and low levels of lamin A/C and B1 are reflected on the right, where the contrast range was varied for visibility. **B**) For the HT1080 cells, lamin A/C fluorescent intensities were plotted against lamin B1 and fitted with a linear trendline. The trendline has a Pearson coefficient of 0.474 and a slope of 0.248. The data points are additionally colour-coded for the corresponding nuclear solidity. On the right, representative images from HT1080 nuclei expressing high or low levels of lamin A/C and B1 are depicted with different contrast ranges for visibility. **C**) For MDA-MB-231, the lamin A/C fluorescent intensities plotted against lamin B1 were fitted with a linear trendline. The trendline has a Pearson coefficient of 0.336 and a slope of 0.167, indicating stoichiometric decoupling, in which the expression of one lamin no longer predicts the presence of the other. The data points are additionally colour-coded for the corresponding nuclear solidity. On the right, representative images from MDA-MB-231 nuclei expressing high and low levels of lamin A/C and B1 are depicted with different contrast ranges for visibility. The number of nuclei imaged is n = 70 for HeLa cells, n = 82 for HT1080 cells and n = 64 for MDA-MB-231 cells. Scale bars represent 10 μm. Imaged over three consecutive experiments.

### Lamin A/C, but not lamin B1, correlates with increased nuclear solidity

Figure 2 suggested that highly deformed nuclei, as indicated by lower solidity values, were found for lower expression levels of both lamin A/C and B1 types. To investigate the individual contribution of lamin expression on nuclear abnormalities, we further performed single-cell correlation analysis of lamin A/C and B1 levels with the corresponding nuclear solidity (Figure 3A-F). Linear fits of the data showed that higher expression levels of lamin A/C correlated with increased nuclear solidity, as a statistically significant positive correlation between lamin A/C expression levels and nuclear solidity was found for all cell types (p-value < 0.05) (Table 2). Nevertheless, the fitted slopes differed across cell types. Where HeLa and HT1080 both had the same slope value (2.1*10^−4^), MDA-MB-231 contained a four times higher slope (8.4*10^−4^) and a higher Pearson coefficient. The differences in fitting parameters suggest a stronger association between lamin A/C levels and nuclear shape in MDA-MB-231 cells than in our other cancer cell types (Figure 3 A, B, C).

**Table 2:** Statistical parameters of nuclear solidity and lamin A/C across cell lines.

|  |  |  |  |  |
| --- | --- | --- | --- | --- |
| Cell types: | Slope<br>(Nuc.Sol.:A/C) | Pearson<br>coefficient $r$ | Offset (y-axis intercept) | p-value |
| HeLa | $2.1 \times 10^{-4}$ | 0.266 | 0.967 | $p < 0.05$ |
| HT1080 | $2.1 \times 10^{-4}$ | 0.288 | 0.962 | $p < 0.01$ |
| MDA-MB-231 | $8.4 \times 10^{-4}$ | 0.435 | 0.901 | $p < 0.001$ |

**Figure 3.**
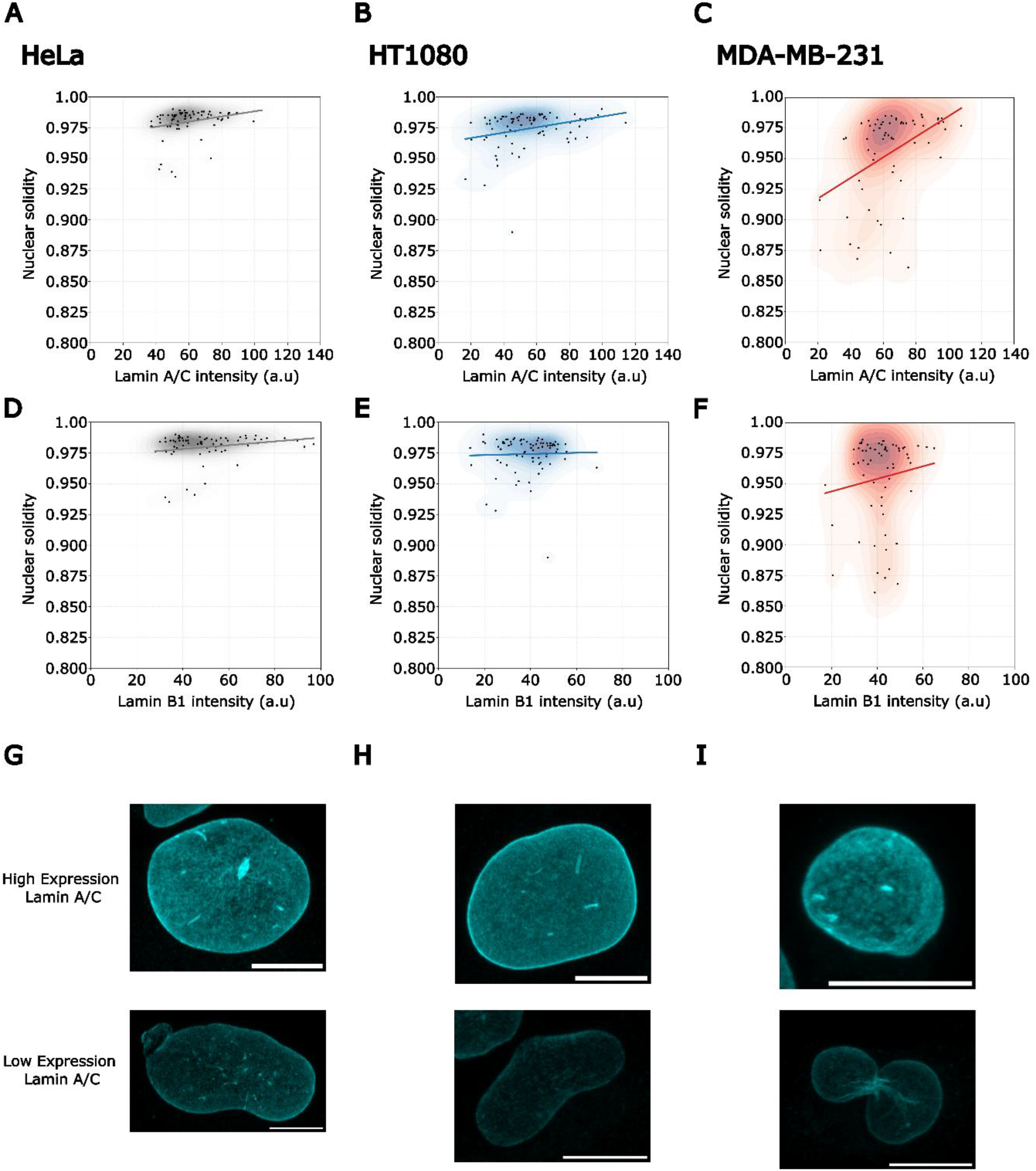
Nuclear solidity is governed by the cell-specific lamin A/C expression levels. **A, D**) HeLa nuclear solidity plotted against lamin A/C, B1 and fitted with a linear trendline. **G**) HeLa nuclei with high and low expression of lamin A/C were depicted. **B, E**) HT1080, nuclear solidity plotted against lamin A/C, B1 and fitted with a linear trendline. **H**) HT1080 nuclei with high and low expression of lamin A/C were depicted. **C-F**) MDA-MB-231, nuclear solidity plotted against lamin A/C, B1 and fitted with a linear trendline. **I**) MDA-MB-231 nuclei with high and low expression of lamin A/C were depicted. All trendlines plotted against lamin A/C and nuclear solidity are significant (p-value < 0.05). Trendlines for lamin B1 and nuclear solidity are not significant (p-value > 0.05). The number of nuclei imaged is n = 70 for HeLa cells, n = 82 for HT1080 cells and n = 64 for MDA-MB-231 cells. Imaged over three consecutive experiments. Scale bars in **G, F** and **I** represent 10 μm.

On the contrary, expression levels of lamin B1 did not correlate significantly with nuclear solidity for any of the human cell lines tested (p-values > 0.05, Table 3). These data, in combination with the data for lamin A/C, revealed that lamin A/C is the main regulative component of the nuclear roundness. Lamin B1, on the other hand, seemed to fulfill a more passive effect on nuclear solidity. NIH3T3 cells were also considered, but the results were inconclusive due to the low lamin B1 expression (Sup. Fig. 2).

**Table 3:** Statistical parameters of nuclear solidity and lamin B1 across cell lines.

|  |  |  |  |  |
| --- | --- | --- | --- | --- |
| Cell types: | Slope<br>(Nuc.Sol.:B1) | Pearson<br>coefficient $r$ | Offset (y-axis intercept) | p-value |
| HeLa | $1.6 \times 10^{-4}$ | 0.207 | 0.972 | $p > 0.05$ |
| HT1080 | $5.0 \times 10^{-5}$ | 0.032 | 0.972 | $p > 0.05$ |
| MDA-MB-231 | $5.2 \times 10^{-4}$ | 0.132 | 0.933 | $p > 0.05$ |

Furthermore, the trendlines each showed a different interception value at the solidity axis, suggesting that the baseline solidity varied per cell type (Table 2, 3). HeLa cells showed the highest baseline solidity, resulting in the smoothest nuclei, followed by HT1080 and the MDA-MB-231 cells respectively. It remained remarkable that the MDA-MB-231 had the lowest baseline solidity and the highest effect of lamin A/C on the nuclear solidity.

Consequently, we hypothesized that there might be a dominant contribution of either lamin A or lamin C on nuclear solidity, which was indistinguishable in this data, and aimed to investigate this further.

### Lamin A and C stoichiometries and interaction strength are cell type specific

Our fluorescence microscopy assays could not distinguish the cell-specific lamin A and lamin C expression levels since our antibodies did not distinguish between the two splice variants. To obtain insight into whether the nuclear shape phenotype is dominated by lamin A or lamin C expression in our cell types, we performed a Western Blot assays to determine the lamin A-to-lamin C ratio. We found that HeLa, HT1080, and MDA-MB-231 all contained monomeric lamin A and lamin C, with HeLa cells having the highest lamin A-to-lamin C ratio, followed by MDA-MB-231 and HT1080 (Figures 4A, B). On the contrary, NIH3T3 did not show any monomeric lamin A and lamin C. The absence of lamin A and lamin C monomers in the NIH3T3 cell lysate, together with our observation that they contained the largest pool of lamin A/C during the population-averaging experiment, led us to further investigate our mild-denaturing Western Blots. Strikingly, we found that the cell lysates of all our cell types contained lamin populations with twice the molecular weight of monomeric lamin A and lamin C, which presumably reflect homodimers (Figure 4C). The ratios of lamin dimers to monomers differed significantly among each cell type, with NIH3T3 showing the highest ratio of lamin dimers to monomers, followed by HT1080, HeLa and MDA-MB-231 cells, respectively.

**Figure 4.**
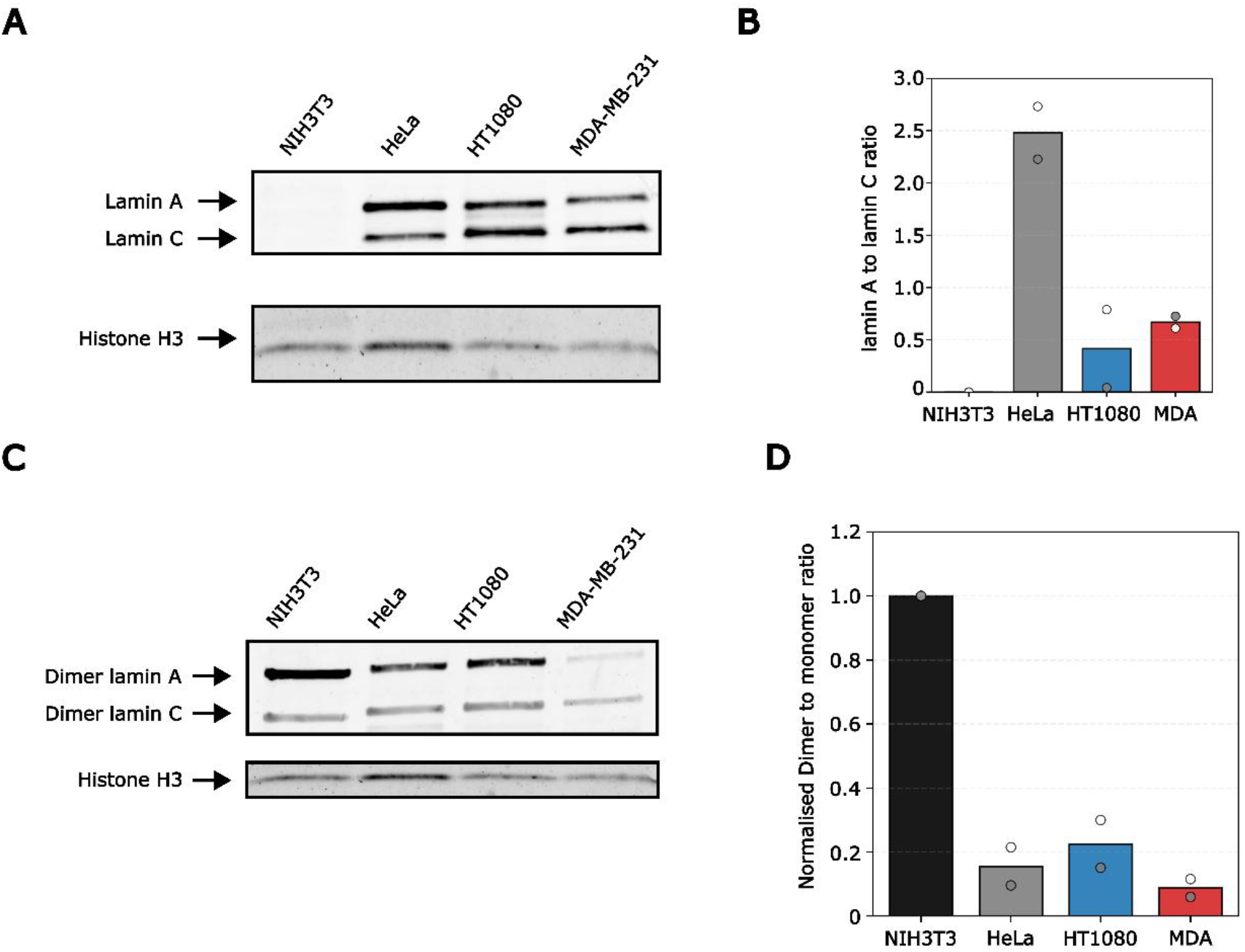
Stoichiometry and dimerisation states of A-type lamins vary per cell type. **A)**Western Blot of soluble whole cell lysate was stained for lamin A/C and histone H3 as a loading control. NIH3T3 showed no monomeric lamin A or C. HeLa, HT1080, and MDA-MB-231 showed monomeric lamin A and C. HeLa contain more lamin A compared to lamin C, HT1080 contain more lamin C compared to lamin A. MDA-MB-231 contains as much lamin A as lamin C. **B)** Quantification of a Duplo Western Blot wherein the ratio of monomeric lamin A to lamin C was plotted. They were first normalised against the loading control. In Western blot analysis, HeLa has the highest A-to-C ratio, followed by MDA and HT1080. **C)** Western Blot of soluble whole cell lysate was stained for lamin A/C and histone H3 as a loading control. The same denaturing and reducing agents used across cell lines produce significantly altered dimer ratios. Dimers of lamin A and lamin C have been observed with equal lamin C dimers in NIH3T3, HeLa, HT1080 and MDA-MB-231. However, Lamin A/C dimers differed significantly among cell lines, with NIH3T3 having the highest levels, followed by HT1080 and HeLa, which showed minimal differences. Minimal lamin A dimers were found in MDA-MB-231. **D)** Quantification of the Western Blot wherein the ratio of lamin A/C dimers to monomers is plotted. Values were internally normalised. NIH3T3 cells have the highest dimer-to-monomer ratio of lamin A/C of 1. HeLa and HT1080 cells differ slightly 0.15 to 0.22. MDA-MB-231 shows the lowest lamin A/C dimer-to-monomer ratio, 0.08. The experiment in **B-D)** was performed twice, grey points represent the first Western blot (Figure S3), white points the second Western blot (Figure S4). Quantifications of the Western blots are provided in the supplementary information (Figure S6).

Interestingly, the presence of lamin A/C dimers correlated well with our observations that MDA-MB-231 showed significantly less solidity than HeLa and HT1080, and even less in NIH3T3 cells. This observation suggests a contributing role of lamin A/C interaction strength on the nuclear mechanical phenotype. However, dimerisation efficiency does not fully account for nuclear morphology, as HT1080 showed a slightly higher dimer ratio than HeLa cells while displaying reduced nuclear solidity.

We hypothesised that the cell-type-specific variation in lamin A and C dimer stability might be caused by cell-type-specific post-translational modifications. To further test this, we performed a Western Blot analysis to identify phosphorylation of lamins (Sup. Fig. 5). Phosphorylation of the nuclear lamina is known to facilitate nuclear envelope breakdown during mitosis and is associated with metastasis and more deformed nuclei, and, as such, might weaken lamin A interactions.^25,26^ We found that only lamin A dimers were phosphorylated, with NIH3T3 showing the highest phosphorylation, followed by HeLa and HT1080. MDA-MB-231 had the fewest phosphorylated dimers. However, variation in the lamin A phosphorylation state did not clarify whether less-stable dimers were present in MDA-MB-231 cells. As such, we speculated that other post-translational modifications are more important for stabilising lamin dimer interactions, but left it to future research to investigate this further. ^27,28^

## Discussion

In this study, we set out to investigate the supportive role of lamin A/C and lamin B1 on nuclear shape. We have particularly focused on cancer cells, including two cell types that are known to be very malignant, since their nuclear shape abnormalities have been correlated with more severe disease prognosis. Using wild-type cell lines, rather than knockouts or overexpression systems, allowed us to investigate the roles of physiologically relevant protein levels. Utilising fluorescence intensities of immunostained lamin A/C and B1 cells as a proxy for lamin A/C and B1 expression levels allowed for single-cell correlation of protein expression levels and nuclear shape phenotyping, indicated by reduced solidity.

Interestingly, we found a positive correlation between lamin B1 and lamin A/C expression for all cell types, suggesting that the different lamin types co-express. In the more invasive cancer cell lines MDA-MB-231 and HT1080, the co-expression analysis showed a shift towards increased lamin A/C expression relative to lamin B1, indicating a more lamin A/C dominated nuclear shape phenotype. Further, single-cell correlation analysis of lamin A/C and B1 with nuclear solidity unravelled that only lamin A/C significantly contributed to nuclear solidity (p-value < 0.05), whereas lamin B1 did not (p-value > 0.05). As such, we presented evidence that the deformability of the nucleus varies across cancer cell types and is facilitated by alterations in the expression levels of lamin A/C, while lamin B1 was not found to influence nuclear deformability, as was previously reported.^13^ Additionally, our observations and measurements in the lamin B1 deficient NIH3T3 cells further highlight the contributing role of lamin A/C to nuclear solidity, as they showed very rounded nuclei while expressing lamin A/C primarily. Though, we cannot exclude cell-type specific additional contributions of chromatin and cytoskeletal influences on nuclear shape. ^29–32^

Previous studies indicated that lamin A is more critical for resisting nuclear deformations due to its stronger binding to the nuclear envelope, while lamin C acts as a less stable, more soluble component of the lamina.^33,34^ To further dissever whether dominant expression of lamin A or C contributes to nuclear shape in our cells, we performed Western Blot analysis and found significant differences in lamin A to lamin C monomer ratios per cell type, which is in agreement with previous observations.^35^ Unsatisfyingly, those findings poorly correlated with the expression levels determined by fluorescence microscopy, as no monomers were detected in NIH3T3 cells, while they expressed the highest level of lamin A/C.

Interestingly, further investigation led us to discover the presence of lamin A and lamin C dimers within all our cell types. The presence of dimers was unexpected since we used protein denaturing conditions, i.e. SDS running buffer in combination with reducing agents, within our Western Blot assays. Though, stable lamin dimers have been previously observed even in 6 M urea.^27^ Most lamin A/C dimers were found in NIH3T3 cells and their presence gradually decreased in HT1080 and HeLa, and was lowest in MDA-MB-231. Since this trend in dimer levels correlated well with the decrease in nuclear solidity, we suggest that reduced lamin A/C dimer levels could facilitate more flexible and abnormal nuclei, as observed in MDA-MB-231 cells. This hypothesis is further supported by the known migration mechanism of HT1080 and MDA-MB-231, where HT1080 cells move through confined spaces by breaking the lamin network and repairing those breaks with blebs that are formed by soluble lamin C.^20,33,36^ MDA-MB-231 cell nuclei deform during migration, which is possibly facilitated by the more flexible lamin network as a consequence of weaker lamin A/C interactions.^20^

Lamin proteins possess exceptional metabolic stability, with turnover lifetimes spanning weeks to months, making remodelling of the nuclear lamina by reduction of total protein content impossible.^37,38^ We hypothesise that tuning of lamin A/C interaction strength, potentially mediated by post-translational modifications within the coiled-coil domains, alters dimerisation state and thereby adapts nuclear lamina rigidity.^28^ This is further supported by identified mutations in the coiled-coil regions, which destabilise lamin A/C dimerisation and cause misshapen nuclei.^39,40^ Future research should further explore the molecular mechanisms underlying lamin A and C interactions and homodimer stabilisation. Our results underline the complexity of nuclear envelope deformations and maintenance. Whereas we show a clear dependence of nuclear shape on lamin A/C expression, our results also show a variability among cancer cells and cell lines that presumably reflects differences in lamin A/C interaction stability. Nevertheless, the enhanced lamin A/C dimer levels in cancer cells with more deformed nuclei may provide a basis for future studies on biomarkers of metastatic progression and potential therapeutic strategies.

## Acknowledgements

We would like to thank Boyd Peters for the fruitful discussions during the research endeavours and Aleksandra Placzek for enabling cell culture lab work. We would like to thank Johan Slotman and Martijn de Gruijter for their help in the microscopy imaging facilities at the Erasmus Medical Centre.

## Funding details

This work was supported by TU Delft internal research funding.

## Disclosure statement

No potential conflict of interest.

## Data availability statement

All data can be asked upon request.

## Author Contributions

MH and HG were involved in the conception and design of this study and wrote the original draft of this paper. MH and AM performed experiments and analyzed and interpreted the obtained data. HG supervised the research. All authors critically revised the intellectual content, approved the final version of this manuscript and agreed to be accountable for all aspects of this work.

## Materials and Methods

### Cell culture

HeLa (ACC 57, DSMZ), HT1080 (ACC 315, DSMZ), MDA MB 231 (ACC 732, DSMZ) and NIH3T3 (ACC 59, DSMZ) were cultured in Dulbecco’s Modified Eagle Medium (11960-044, Gibco) supplemented with 10% (v/v) fetal bovine serum (FBS, Avantor), 1% (v/v) 100x Glutamax (35050-038, Gibco) and 1% (v/v) penicillin/streptomycin. Cells were maintained in a humidified, gas-regulated incubator at 37 °C and 10% CO2. When cells reached 90% confluency, they were passaged at dilutions determined by their growth rate.

### Fixation and Immunostaining

Cells were seeded on 18 mm #1.5H coverslips (0117580, Marienfeld) 24 hours before fixation. Before fixation, DMEM was aspirated, and cells were washed once with prewarmed phosphate-buffered saline (PBS, 20012-019, Gibco). PBS was aspirated, and -20 °C absolute ethanol was added to the cells, which were incubated for 10 minutes at room temperature. Subsequently, cells were washed three times for five minutes with PBS on an orbital shaker (100 RPM). Cells were permeabilised with permeabilisation buffer (PBS, 0.2% (v/v) Triton X-100) for 5 minutes on an orbital shaker (100 RPM). Afterwards, cells were blocked with blocking buffer (PBS, 5% (v/v) Goat serum, ab7481, Abcam) and incubated for 1 hour on an orbital shaker (100 RPM). Blocked cells were stained with mouse anti-Lamin A/C (sc-376248, Santa Cruz) and rabbit anti-Lamin B1 (12987-1-AP, Proteintech) antibodies diluted 1:200 in staining buffer (PBS, 1% (w/v) Bovine serum albumin (BSA, A2153-100G, Sigma-Aldrich)) and incubated for 1 hour at room temperature. After the first staining, the cells were washed three times for 5 minutes with PBS on the orbital shaker (100 RPM). Secondary antibodies, goat anti-mouse Alexa Fluor 488 (AB150113, Abcam) and goat anti-rabbit Cy3b (SA00009-2, Proteintech), were diluted 1:400 in staining buffer and incubated with cells for 1 hour at room temperature. Stained cells were washed three times for 5 minutes with PBS on the orbital shaker (100 RPM). Post-fixation of cells was performed with 2% PFA for 10 minutes at room temperature, followed by three washes with PBS for 5 minutes each.

### Data acquisition

Cells were imaged on the Leica SP8 with a 63 × 1.4 NA oil objective. Z-stacks of entire nuclei were acquired with z-steps of 160 nm and xy pixel sizes of 80 nm, optimised for Nyquist sampling while minimising photobleaching. Lamin A/C stained with Alexa Fluor 488 was illuminated with the 488 nm laser, and Lamin B1 stained with Cy3b was illuminated with 561 nm. To avoid signal saturation, the laser power and gain were adjusted to ensure that all pixel values fell within the detector’s dynamic range. These settings were kept constant over all measurements.

### Data processing

All z-stacks are processed using ImageJ. Z-stacks were converted to maximum intensity projections (MIP). The projected nuclei were used to generate binary masks from the lamin A/C channel using ImageJ’s default auto-thresholding to define the nuclear region of interest. The same binary mask was used to measure nuclear solidity. Nuclear solidity is calculated by dividing the measured fluorescent area by the area of the convex hull. The lamin A/C binary mask was used to measure the mean grey values of lamin A/C and B1. This was done to ensure that blebs that did not contain lamin B1 were still accounted for. Correlation of lamin A/C or B1 fluorescence on nuclear solidity was determined by a linear regression. Mechanical efficacy was defined as the slope of the trendline (nuclear solidity/fluorescent intensity). Statistical significance was subsequently determined, with p-values < 0.05 considered significant. For each cell line, approximately 20 nuclei were analysed across each consecutive experiment to ensure reproducibility.

### Western Blot

Cells were grown in 10 cm tissue culture plates until confluency and harvested. Subsequently, the cell suspension was centrifuged for 10 minutes at 300xg, and the supernatant was aspirated. The cell pellet was afterwards frozen at -20 until further use. Before lysation, the cell pellet was washed once with phosphate-buffered saline (PBS) and again centrifuged for 10 minutes at 300xg. Lysis of whole-cell pellets was performed by adding 300 μL 1x Radioimmunoprecipitation Assay (RIPA) buffer and incubating for 15 minutes on ice. Lysates were centrifuged to collect the soluble fraction. The soluble fraction was diluted 1:1 with 2x Laemmli buffer supplemented with 5% β-mercaptoethanol and heated at 85 °C for 10 minutes. 30 μL of the boiled sample was loaded onto an SDS-PAGE gel and run for approximately 1 hour. The separated proteins were blotted onto nitrocellulose using a Trans-blot Turbo (Bio-Rad). The blot was blocked with 3% milk blocking buffer for 1 hour at room temperature. The phosphate protein detection Western Blot was blocked with 1% BSA. Subsequently, the primary antibody was incubated overnight at 4 °C (Table 4). After primary antibody incubation, the sample was washed three times for five minutes each in PBS supplemented with 0.1% Tween-20. Following, the incubation of the secondary antibody for 1 hour at room temperature (Table 4). After incubation, it was washed three times with PBST, then imaged on the Odyssey Clx (LI-COR Bioscience).

**Table 4:** List of Antibodies used for Western Blot.

| **Table 4: List of Antibodies used for Western Blot** |  |  |  |  |  |
| --- | --- | --- | --- | --- | --- |
| Protein target | Dilution | Dye | Manufacture | Host | Catalog number |
| Lamin A/C | 1:6000 | - | Proteintech | Rabbit | 10298-1-AP |
| Histone H3 | 1:3000 | - | Proteintech | Rabbit | 17168-1-AP |
| Phospho-Lamin A/C (Ser22) | 1:1000 | - | Cell signaling | Rabbit | 13448T |
| Anti-Rabbit | 1:5000 | IRDye 800cw | LI-COR Bioscience | Goat | 926-32211 |

## Notes

### Competing Interest Statement

The authors have declared no competing interest.

### Summary of Updates

Figures 3 and 4 were not depicted correctly in the original upload.

