## Supplementary material for "Expression levels and dimer abundance of lamin A/C direct nuclear shape integrity in malignant cancer cells": Supplemenatry Information, Figures S1-S6 (from another version of this preprint: version 1, 2026-05-13)

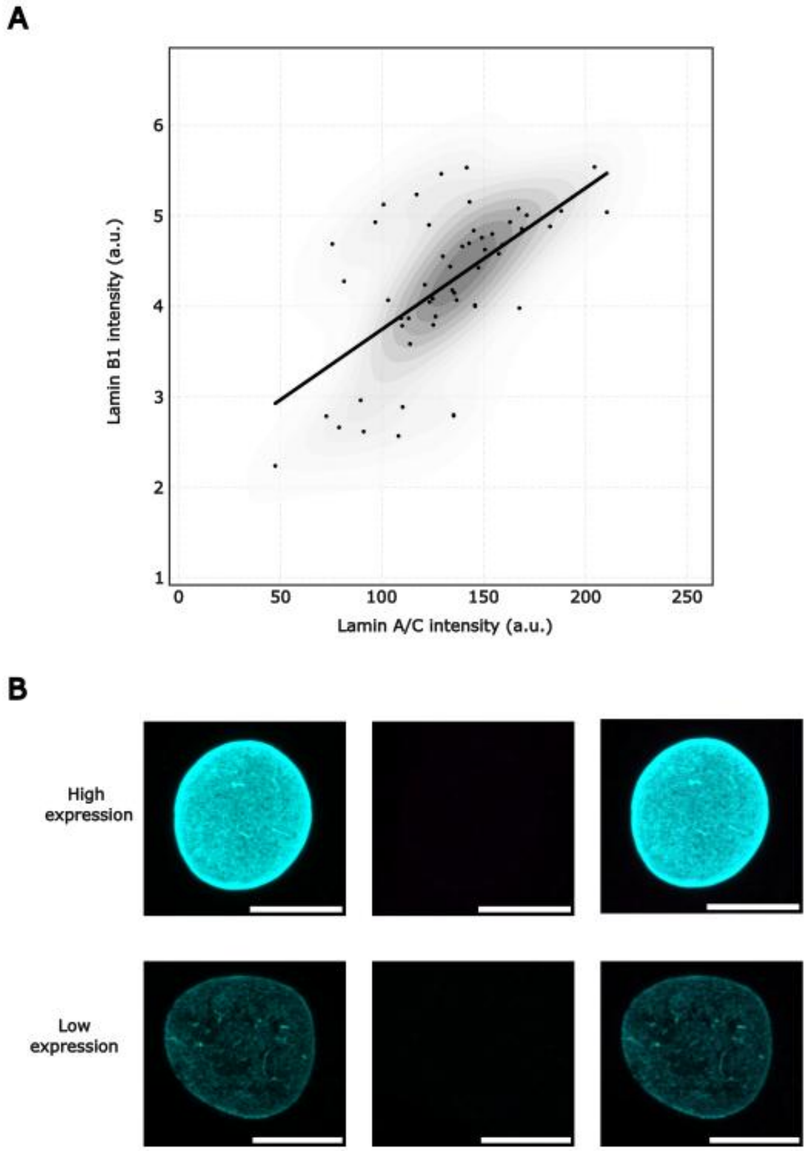


Figure S1: **NIH3T3 lamin A/C to lamin B1 ratios**
**A**) NIH3T3 fluorescent intensities of lamin A/C and B1 are plotted against each other and fit with a linear trendline. The trendline has a Pearson coefficient of 0.616 and a slope of 0.016. The lamin B1 signal is close to the background signal. **B)** Representative images of NIH3t3 nuclei expressing high and low levels of lamin A/C and B1. The number of nuclei imaged is n = 54. Scale bars represent 10 µm.

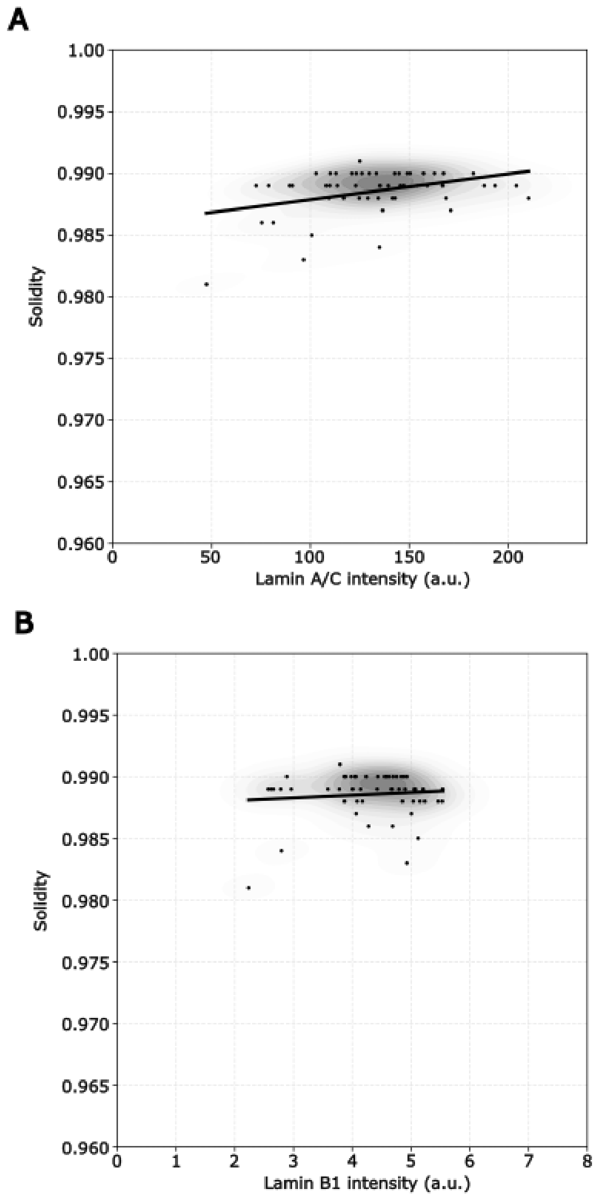


Figure S2: **Effects of lamin A/C and lamin B1 expressions on nuclear solidity of NIH3T3**
**A**) NIH3T3 nuclear solidity against lamin A/C scatterplot, fitted with a linear trendline with a Pearson correlation of 0.375 (p-value 0.005). **B**) NIH3T3 nuclear solidity against lamin B1 scatterplot, fitted with a linear trendline with a Pearson correlation of 0.102 (p-value 0.5).

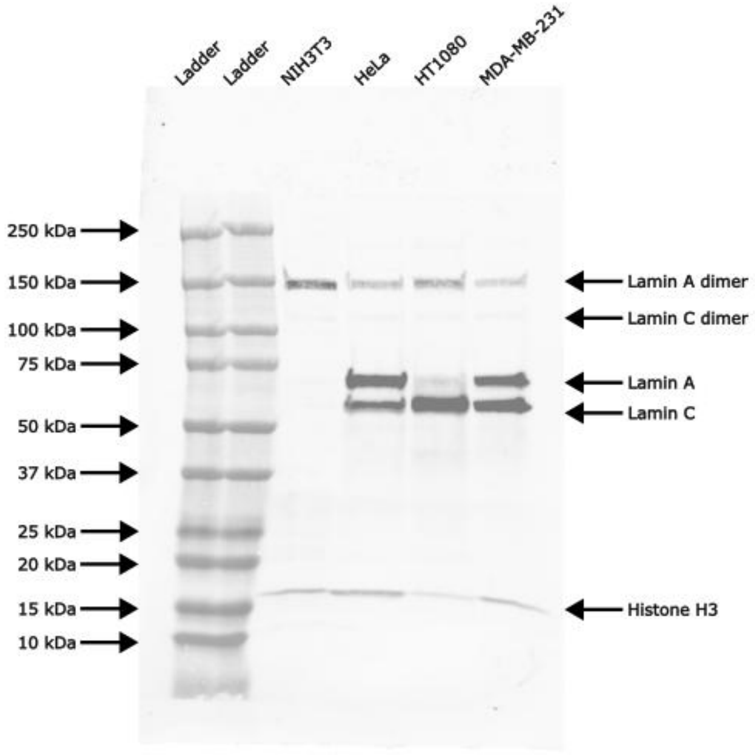


Figure S3: **Western Blot for identification of the ratio of lamin A to lamin C and dimers in cell lines**
Western Blot used to determine the ratio of lamin A to lamin C and their corresponding dimers, of NIH3T3, HeLa, HT1080 and MDA-MB-231. NIH3T3 did not contain any monomeric lamin A and C. However, it did contain the most dimers of lamin A and lamin C. HeLa, HT1080 and MDA-MB-231, contained dimers and monomers of lamin A and lamin C. The loading control used is histone H3.

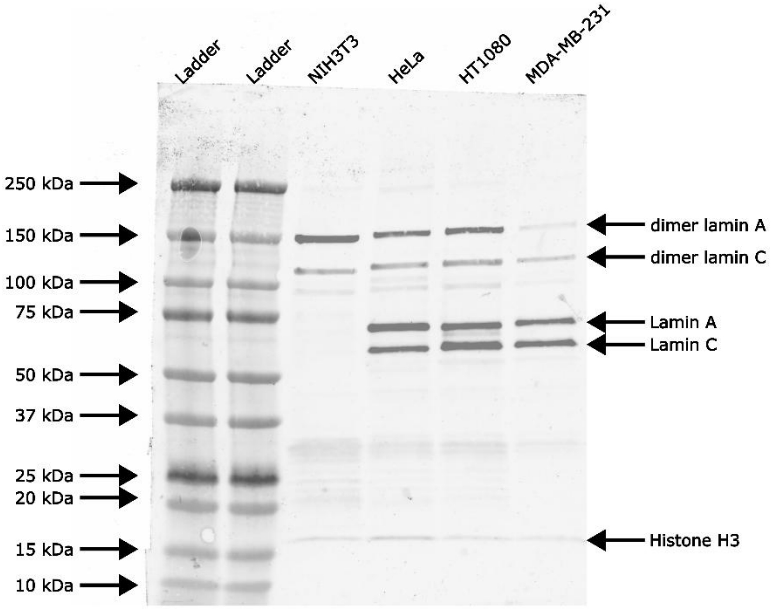


Figure S4: **Western Blot for identification of the ratio of lamin A to lamin C and dimers in cell lines**

Western Blot used to determine the ratio of lamin A to lamin C and their corresponding dimers, of NIH3T3, HeLa, HT1080 and MDA-MB-231. As previously observed, NIH3T3 cells did not contain any monomeric lamin A or C. However, it contained the most dimers of lamin A and lamin C. HeLa, HT1080, and MDA-MB-231 contained dimers and monomers of lamin A and lamin C. The loading control used is histone H3.

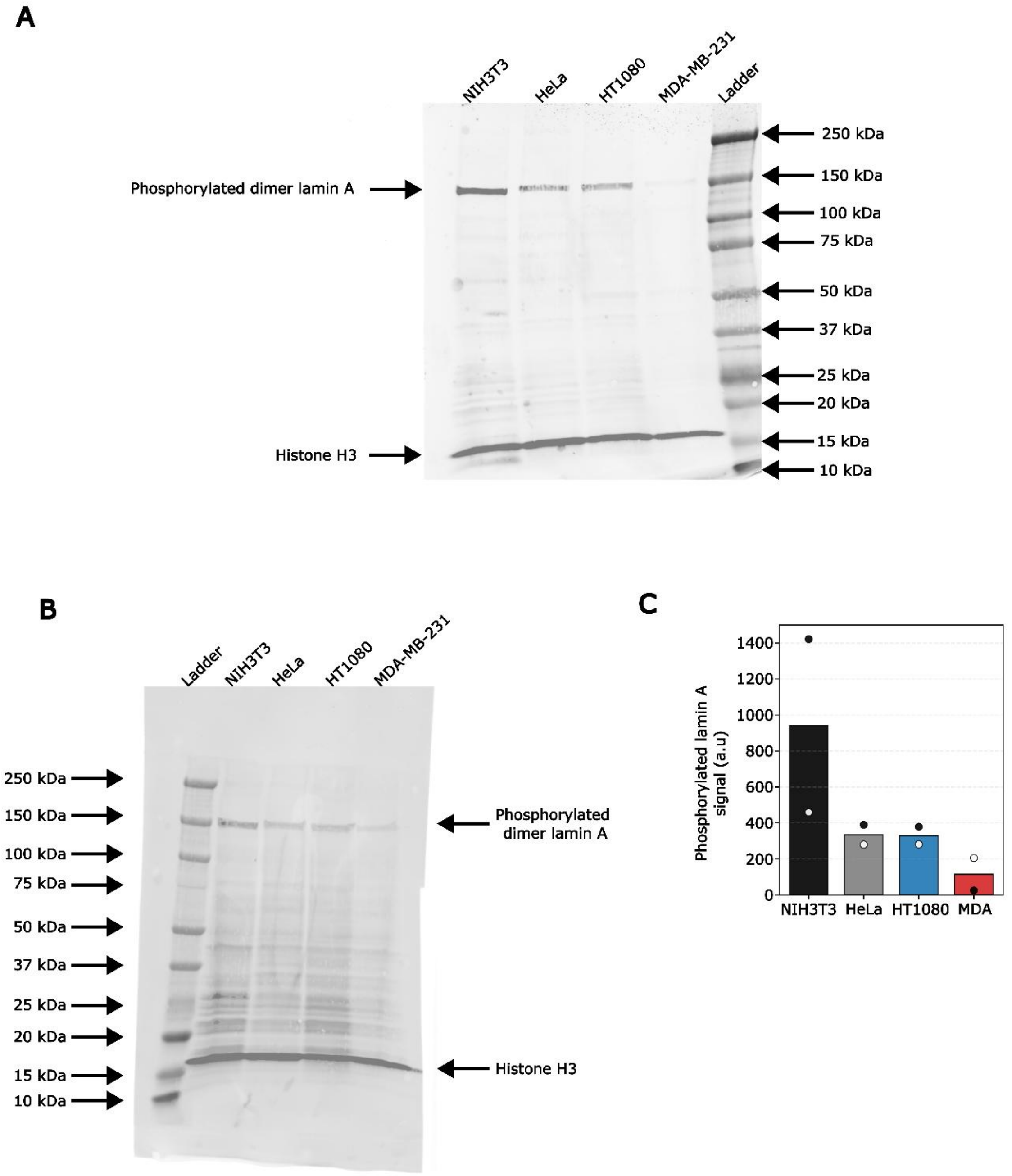


Figure S5: **Western Blot for identification of lamin A/C phosphorylation of Serine 22**

**A**) Western Blot used to determine the post-translational modification phosphorylation of Serine 22 of lamin A/C. NIH3T3 shows the most phosphorylated lamin A dimers, followed by HT1080, HeLa and MDA-MB-231, which contain the least phosphorylated lamin A dimers. Histone H3 was used as a loading control. **B**) Western Blot used to determine the post-translational modification phosphorylation of Serine 22 of lamin A/C. NIH3T3 shows the most phosphorylated lamin A dimers, followed by HT1080, HeLa and MDA-MB-231, which contain the least phosphorylated lamin A dimers. Histone H3 was used as a loading control. **C**) Phosphorylated lamin A dimer signal in arbitrary units. Showing that MDA-MB-231 contains the least amount of phosphorylated lamin A. White points represent the westernblot in **A**, black points represent the points in **B**.

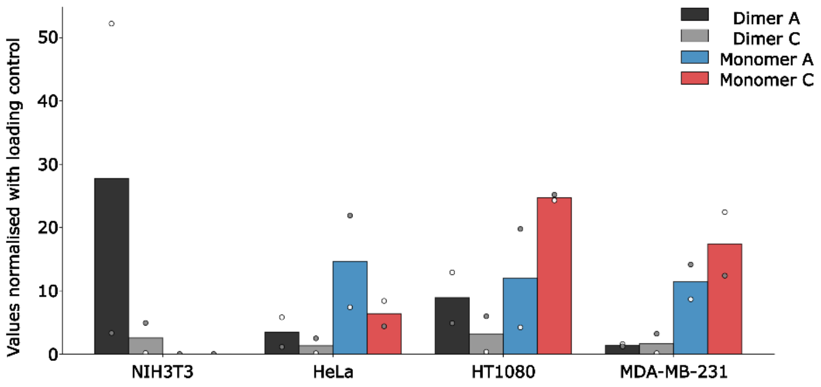


Figure S6: **Quantitative Western Blot analysis of Lamin A and C oligomerisation states**

Western Blot in Figures S3 and S4 was used to quantify monomeric and dimeric lamin A and C. Each point represents the value from the corresponding Western Blot, normalised to the loading control. NIH3T3 exhibits no monomers, while MDA-MB-231 exhibit almost no dimers. White points represent Western blot in Figure S3, Gray points represent Western blot in Figure S4.
